# Conjugation dynamics and persistence of a carbapenem resistance gene *bla*_OXA-72_ from *Acinetobacter pittii* to *Acinetobacter baumannii*

**DOI:** 10.64898/2026.06.25.734492

**Authors:** Kenneth Bongulto, Hisamichi Tauchi, Satoru Suzuki, Kozo Watanabe

**Author notes:** ADDRESS CORRESPONDENCE TO: Kozo Watanabe.

## Abstract

Carbapenem-resistant *Acinetobacter* (CRA) has been associated with increased morbidity and mortality in clinical settings. In this study, we explored the transfer potential of a mobilizable plasmid-harboring *bla*_OXA-72_ gene between *Acinetobacter* species originating from patient, municipal wastewater, and pig farm wastewater. PCR-based evidence suggested putative transfer of *bla*_OXA-72_ from *Acinetobacter pittii* to *Acinetobacter baumannii.* In this pair, the apparent frequency of PCR-marker-positive putative transconjugants varied depending on temperature and meropenem supplementation, with higher number observed at 27°C compared to 17°C and 37°C. Likewise, the presence of antibiotic pressure yields to higher apparent conjugation frequency, however this observation was limited to a singled donor-recipient pair. Further, we revealed a phenotypic conversion in terms of meropenem susceptibility and a fitness cost in the putative transconjugants. While whole genome sequencing did not conclusively verify the presence of *bla*_OXA-72_ or fully resolved plasmid configuration, Oxford Nanopore read mapping consistently detected the chromosomal *strA* gene in all isolates. In contrast, only a limited number of reads aligned with *bla*_OXA-72_ gene, *traC*, or the complete plasmid sequences. Comparative analyses further revealed variations in the surface-associated factors and defense systems composition of the recipient strains, which could be considered as barriers in conjugation. Lastly, the persistence of PCR-detectable marker genes in putative transconjugants was variable and generally unstable over a 30-day period. Overall, these findings provide preliminary insights into the factors that may influence horizontal gene transfer and short-term maintenance of *bla*_OXA-72_.

**IMPORTANCE:** Carbapenem-resistant *Acinetobacter* (CRA) is a serious clinical concern linked to elevated morbidity and mortality. This study shed light on how the carbapenemase gene *bla*OXA_72 spreads in the environment via horizontal gene transfer, Using isolates of *Acinetobacter pittii* and *Acinetobacter baumannii* from municipal wastewater treatment plant, we show that conjugation efficiency depends on temperature and antibiotic exposure. The process also incurs fitness costs, and the transferred gene persists only briefly in recipient cells. These results emphasize the importance of environmental sources as reservoirs for antimicrobial resistance gene exchange. Additionally, variations in surface structures and defense systems may act as potential barriers to conjugation. Overall, the work improves our understanding of AMR movement and supports the need for better integrated surveillance under a One Health approach.

## INTRODUCTION

In recent years, the emergence of multidrug resistant *Acinetobacter* species has been reported to cause infections and mortality in healthy and immunocompromised individuals. The enrichment of antimicrobial resistance genes (ARGs) in plasmids has greatly influenced the development of resistance in *Acinetobacter* species (1, 2). Additionally, this phenomenon allowed the horizontal transfer of ARGs from one bacterial species to another which has facilitated the adaptation of environmental *Acinetobacter* species to nosocomial settings. Most importantly, carbapenem resistance genes which confer resistance to last resort antibiotics have been found enriched in plasmid replicons of *Acinetobacter* species (3). The mechanism of carbapenem resistance involves the production of carbapenem-hydrolyzing class D β-lactamases (4). This encompasses the intrinsic *bla*_OXA-51-like_ gene and the horizontally acquired genes *bla*_OXA-23-like_, *bla*_OXA-24/40-like_, and *bla*_OXA-58-like_ genes (5, 6). One of the most well-known carbapenemases is the *bla*_OXA-72_ gene, a variant of the *bla*_OXA-24/40-like_ β-lactamases, which is generally plasmid-borne in nature (7–9).

Although initially identified in *Acinetobacter baumannii*, *bla*_OXA-24/40-like_ β-lactamases such as *bla*_OXA-72_ gene were less commonly found in carbapenem-resistant *Acinetobacter* species compared to other β-lactamases (10, 11). In clinical settings, carbapenem treatment, ICU stay, and mechanical ventilation were some of the common risk factors for carriage of *bla*_OXA-72_-producing *A. baumannii* (12). While known to be associated with clinical *Acinetobacter* isolates (13, 14), there were few reports of *bla*_OXA-72_ in *Acinetobacter* from the environment (15, 16) and animal sources (17, 18). Interestingly, the GC content of *bla*_OXA-24/40-like_ β-lactamases was considerably lower than the average for *A. baumannii* genome which indicates that these genes originated outside this species (19). Recently, studies evaluated the transfer potential of plasmid harboring *bla*_OXA-72_ between *Acinetobacter* species but showed different conjugation outcomes (20, 15). Despite increasing reports of *bla*_OXA-72_–harboring plasmids in *Acinetobacter* spp., key aspects of their dissemination remain unresolved. The efficiency of interspecies transfer across strains originating from distinct ecological niches including clinical, environmental, and agricultural settings remains unclear, with previous studies reporting inconsistent outcomes.

Conjugation dynamics is governed by two kinetic processes known as the conjugation frequency and growth dynamics. Conjugation frequency or the efficiency of gene transfer is quantified by the ratio of transconjugants to the number of donors or recipients (21). Conjugation frequency is reported to be influenced by environmental factors such as temperature and antibiotic pressure. Temperature variations can impact bacterial growth rates and conjugation efficiency (22–25, 26, 21). Further, presence of antibiotics can generally induce the process of conjugation (27–30). While temperature and antibiotic exposure are known to influence conjugation, their combined effects under environmentally relevant conditions have not been systematically examined, as most studies are conducted at standard laboratory temperatures that may not reflect natural reservoirs. Additionally, conjugation barriers may affect conjugation efficiency. These barriers include the bacteria’s ability to form intact capsule (31) and the presence of bacterial defense systems (32, 33). Given that *Acinetobacter* is a major driver of resistance dissemination, identifying the factors that facilitate conjugation is essential for managing the spread of resistance across diverse ecological settings.

In assessing the dynamics of horizontal gene transfer (HGT), it is important to consider the persistence of newly acquired plasmids and associated resistance genes, as this determines the long-term success of transmission. Acquisition of a *bla*_OXA-72_–harboring plasmid can impose a fitness cost, which may lead to plasmid loss in the absence of selective pressure. The persistence of plasmid-associated ARGs has been reported to range from a few days to several weeks (34, 35). Plasmid loss can occur in the absence of selective pressure, although resistance genes may, in some cases, be maintained through mechanisms such as chromosomal integration (36). Although chromosomal integration of antimicrobial resistance genes is considered relatively rare, it has been reported in certain contexts (37). However, the extent to which these processes contribute to the persistence of *bla*_OXA-72_ in *Acinetobacter* remains poorly characterized.

Together, these gaps highlight the need to better understand the environmental and biological factors shaping the transfer and persistence of *bla*_OXA-72_–harboring plasmids. The study examines HGT dynamics involving *bla*_OXA-72_ between *A. pittii* and *A. baumannii*. We hypothesize that antibiotic pressure and temperature might influence the conjugation frequency in this donor-recipient pair, as well as the persistence of PCR-detectable marker genes in putative transconjugants over time. We therefore investigated how these conditions affected conjugation frequency, as well as the presence of PCR-detectable markers consistent with putative transfer. Additionally, we assessed potential fitness costs in putative transconjugants and the presence of marker genes over time under different temperature and antibiotic conditions.

## 2. MATERIALS AND METHODS

### 2.1. Selection of potential pairs for conjugation

The phenotype and genotype of *Acinetobacter* isolates were considered in choosing the donor and recipient bacteria in the conjugation assay. The donor bacteria must possess a conjugative and mobilizable plasmids, and the plasmid must harbor an ARG. In contrast, the recipient bacteria must possess a chromosomal ARG and lack any conjugative plasmids. The genome profiles of the donor and recipient bacteria were determined using our previous whole genome sequence data from a previous investigation (38). Oxford Nanopore Technology (ONT) and Illumina NovaSeq 6000 were used to obtain hybrid assembled genomes. The donor and recipient bacteria should have distinct antibiograms for selection. To determine the antibiograms of the isolates, the *Acinetobacter* were tested against multiple antibiotics (i.e. aminoglycosides, beta-lactam, carbapenem, cephalosporins, fluoroquinolones, and tetracycline). Antibiograms were carried out following the guidelines outlined by the Clinical Laboratory Standards Institutes (39). Plasmid replicons of the isolates were assessed to determine the presence of conjugative and mobilizable plasmids carrying ARGs in donor cells. Recipient cells were checked for chromosomal ARGs which confer resistance to an antibiotic.

### 2.2. Characterization of the donor and recipient *Acinetobacter* strains

We used one donor *Acinetobacter* strain in the conjugation assay. The donor strain was isolated from the influent of a municipal wastewater treatment plant in Matsuyama, Japan and was identified as *Acinetobacter pittii* (EA01). The isolate possessed two plasmid replicons, one of which possessed a carbapenemase gene *bla*_OXA-72_. Further, EA01 showed resistance to meropenem and was chosen as the donor strain for the conjugation assay. Four *Acinetobacter* species, which include 2 *A. baumannii* strains, 1 *A. towneri*, and 1 *A. johnsonii*, were selected as recipient bacteria for the conjugation assay. The first *A. baumannii* strain (EA06) was isolated from the influent of the same municipal wastewater treatment plant. The isolate possessed 6kb and 8kb plasmid replicons, both of which were non-mobilizable and lacked ARGs. Additionally, a patient-derived *A. baumannii* strain (EC04) harboring 11kb and 44kb plasmid replicons was isolated from a sputum sample. The plasmids of the patient-derived *A. baumannii* strains were both non-mobilizable and lacked ARGs. Both isolates were susceptible to meropenem but were resistant to certain antibiotics (i.e. EA06 was tetracycline resistant, while EC04 was ciprofloxacin resistant). The other recipient bacteria include *A. towneri* (EA13) and *A. johnsonii* (EA18), both isolated from pig wastewater. The isolate EA13 possessed 9 plasmids while EA18 harbored 20 plasmids **(Supplementary Table S1)**.

### 2.3. Bioethical clearance

The study sought approval from the Ehime University Hospital Clinical Research Ethics Review Committee with approval number 2511001. The clinical isolate EC04 used as one of the recipient strains in the conjugation assay was anonymized.

### 2.4. Conjugation assay

Filter-mating was performed to test for putative transconjugant recovery from selected donor and recipients **(Supplementary Fig. S1)**. The donor was cultured in meropenem-supplemented Luria-Bertani (LB) broth (Becton, Dickinson and Company, New Jersey, United States), while recipient bacteria were cultured in LB broth for ∼20 hours at 30°C. The bacterial density for both donor and recipient bacteria was determined to have an optical density of OD_600_ 1.0. The donor and recipient bacteria were centrifuged at 4000 rpm to collect the bacterial pellets. The pellets were washed with 1× phosphate-buffered saline (PBS) solution to remove residual antibiotics. Equal volume (1:1) of donor and recipient cells were mixed in a sterile 1.5 ml Eppendorf tube and were filtered through a 0.22 µm membrane filter (Merck Millipore, Massachusetts, United States) by vacuum filtration. To observe the effect of antibiotic pressure and temperature in conjugation frequency, the membrane filters were aseptically transferred onto an LB agar with and without meropenem (2 µg/ml) supplementation and incubated at different temperatures (17°C, 27°C, 37°C) for ∼20 hours. After incubation, the membrane filters were aseptically placed in a sterile tube with 1ml of 1× phosphate-buffered saline (PBS) solution and vortexed for 1 minute to disrupt the adhering cells. Spread plating was performed using 100 µl of the mixed bacterial cells onto LB agar supplemented with a combination of 4 µg/ml of meropenem (Kanto Chemical Co. Inc., Tokyo, Japan) and 16µg/ml of tetracycline (Nacalai Tesque, Inc., Kyoto, Japan) to select for putative transconjugants. For donor and recipient selection, 1×10^-6^ diluted bacterial suspension were used. LB agar supplemented with 4 µg/ml of meropenem, and 16 ug/ml of tetracycline were used for the selection of donors and recipients, respectively. Additionally, we implemented stringent contamination controls to rule out donor carryover and mixed colonies. Donor-only and recipient-only were plated on the LB agar supplemented with 4 µg/ml of meropenem, and 16 ug/ml of tetracycline at the same time as the mating mixtures. The agar plates were incubated at 30°C for ∼20 hours. The colony forming unit (cfu) was counted for the donor, recipient, and transconjugant plates. Conjugation frequency was determined by getting the proportion of transconjugants (T*cfu*) over the sum of both recipient (R*cfu*) and transconjugant (T*cfu*) bacteria 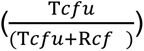.

### 2.5. Species identification and detection of ARGs and plasmid genes in putative transconjugants

To confirm the identity of putative transconjugants and rule out donor contamination, the 16S rRNA gene was amplified using the primers 27F (AGAGTTTGATCCTGGCTCAG) and 1525R (AGAAAG- GAGGTGATCCAGCC) (40). The PCR conditions include 94°C initial denaturation for 5 minutes followed by 30 cycles of 94°C, 60°C, and 72°C. The final extension was done for 7 minutes at 72°C. Further, ARG- and plasmid-specific primers were used to detect presence of ARG and plasmids in the putative transconjugant cell. The presence of the chromosomal ARG *strA* in recipient bacteria EA06 and the plasmid ARG *bla*_OXA72_ in the donor bacteria EA01 were chosen as the marker genes in the putative transconjugants. We designed primers targeting the conjugative plasmid *traC* and the ARG *bla*_OXA-72_ of the donor bacteria EA01 to detect donor- and recipient-specific marker genes in the putative transconjugants. The primers used and PCR conditions for the detection of marker genes in the putative transconjugants were outlined in **Supplementary Table S2**. PCR amplicons were visualized by electrophoresis on a 1.5% agarose gel. Subsequently, the amplified products were purified using the QIAquick PCR Purification Kit (QIAGEN) and sent to Eurofins Genomics Ltd. for Sanger sequencing. The resulting sequence data were analyzed using the NCBI BLAST. Sequence matches were evaluated and prioritized based on E-value, query coverage greater that 90%, and sequence identity exceeding 97%.

### 2.6. Antimicrobial susceptibility testing

To determine whether putative transfer of ARGs influence the putative transconjugants susceptibility against meropenem, broth microdilution method was performed. The antibiogram of the original recipient bacteria and the putative transconjugants against meropenem were determined using the minimum inhibitory concentration (MIC) following the CLSI guidelines in determining the susceptibility of bacterial isolates against antibiotics (39). To determine the phenotype against meropenem, MIC value of ≤ 2 µg/ml was considered susceptible while ≥8 µg/ml was resistant. Meropenem antibiotics were prepared in two-fold concentrations (0.5, 1, 2, 4, 8, 16, 32, 64, and 128 µg/ml) in Mueller-Hinton broth. The bacterial density of each test isolate was adjusted to 0.5 McFarland standard prior to the inoculation in the 96-well plates with different meropenem concentrations. One hundred microliters of the inoculum were used for the broth microdilution assay. The 96-well plates were incubated overnight (∼20 hours) at 30°C. The MIC was determined by using a colorimetric assay by adding resazurin dye to the 96-well plate. Wells without viable cells after exposure to the antibiotics appeared as color blue, while wells with living cells appeared as red. The MIC value is defined as the lowest concentration of antibiotics which prevented the growth of bacteria.

### 2.7. Growth curve for fitness cost

The growth dynamics of both the original recipient (EA06) and putative transconjugants were obtained. Three biological replicates of the putative transconjugant and EA06 were subjected to 24-hour culture in LB broth without antibiotic supplementation. The growth rate for each setup was obtained in triplicates at eight different timepoints: 0, 2, 3, 4, 8, 10, 12, 16, and 24 hours. Bacterial growth was determined by using 600 µl of each liquid culture in a spectrophotometer using optical density at 600 nm (OD_600_). The OD_600_ values of the putative transconjugants were compared with the OD_600_ values of the original recipient bacteria.

### 2.8. Persistence assay for PCR-detectable marker genes and recoverable cells

We propagated putative transconjugants in LB agar supplemented with 4 µg/ml meropenem overnight. A single colony was picked out and inoculated in 10 ml Luria Bertani (LB) broth with 4 µg/ml overnight and were set as day 0. The bacterial cells were cultured every day by transferring 10 µl of inoculum to 10ml LB broth (1:200) for 30 days. The serial passage of putative transconjugants includes three different experimental set-ups. The experimental set-ups include set-up A – without meropenem supplementation, B – with 2 µg/ml meropenem, and C – with 4 µg/ml meropenem. Additionally, all experimental set-ups were exposed to varying temperatures (17°C, 27°C, 37°C) for 30 days. Each set-up was prepared in triplicates. To select for putative transconjugants, spread plating on LB agar supplemented with tetracycline (16 µg/ml) and meropenem (4 µg/ml) was conducted on days 1, 2, 3, 4, 5, 6, 7, 10, 20, and 30. Three single colonies were randomly selected for polymerase chain reaction (PCR) targeting the ARGs (*strA* and *bla*_OXA-72_) and plasmid-specific gene (*traC*) to determine PCR-detectable marker gene persistence (41). The ARGs or plasmids in the putative transconjugant were considered to persist if marker genes were detected in one of the three randomly selected colonies. In the absence of the marker genes, the result will be noted as 0 and will be identified as persisting cells rather than putative transconjugants.

### 2.9. Whole genome sequencing and *de novo* assembly

The whole genome sequencing was performed using the MinION (Oxford Nanopore Technology). Putative transconjugants from days 0, 1, 3, 4, 6, and 30 were subjected to whole genome sequencing to evaluate the presence of ARGs and plasmids and to detect potential genetic changes. Genomic DNAs (∼1 µg) were subjected to library preparation using the Ligation Sequencing Kit V14 (SQK-LSK114). The long-read sequences were generated using the MinION flow cell R10.4.1 (FLO-MIN114) and a MinION MK1B sequencing device (Oxford Nanopore Technology). MinKNOW v25.05.14 software was used for data acquisition. Dorado v1.4.0 was used for basecalling the resulting reads from pod5 files, followed by demultiplexing using the Porechop tool v4.9.1 (42). The sequence quality was assessed using NanoPlot v1.41 (43). Unicycler v0.5.1 with default settings was used to assemble the long-read sequences (44). The assembled genomes were visualized using Bandage v0.9.0 (45). Subsequently, we mapped the Oxford Nanopore reads to reference sequences of *bla*_OXA-72,_ *strA*, *traC*, the mobilzable plasmid pApiEA01a, and the conjugative plasmid pApiEA01b using Minimap2 (46). The alignments were then sorted, indexed, and filtered at MAPQ ≥20 and ≥30 with SAMtools (47).

### 2.10. Bioinformatic analyses

Plasmid sequences were annotated using the default parameter of Bakta (48). The annotated .gbk files were utilized for gene synteny analysis using the geneviewer package in R (49). MOBsuite v.3.1.7 (50) was used to determine the mobility of the plasmid replicons. Plasmid *dif* modules were predicted using the default parameter of pdifFinder (51). The ARGs were detected using the ResFinder v.4.6.0 database (52). DefenseFinder v2.0.1 with default parameter was utilized to predict bacterial defense systems (53). The default parameter for CRISPRCasFinder (54) was used to determine the spacer sequences. The CRISPR-Cas system was further investigated by conducting nucleotide sequence similarity searches with BLASTn (55) using the default parameters optimized for short queries. The individual spacers were used as the query while the plasmid sequence was used as the subject in the alignment. Hits showing ≥90% in sequence identity and few gaps or mismatches were considered. Additionally, the protospacer adjacent motif (PAM) for CRISPR-Cas Type I-F was investigated for presence of CC or NCC immediately upstream (5’) of the protospacer. The capsule (K locus) and outer core oligosaccharide (OC locus) were determined using the KAPTIVE tool v1.3.0 with default settings against the curated *Acinetobacter* K and OCL reference databases (56). Insertion sequence (IS) elements were predicted using the default parameter of MobileElementFinder v1.0.3 (57). To investigate potential mechanism associated with continued growth on meropenem-supplemented media, whole genome sequence of persisting cells recovered from day 30 and day 0 were compared using MUMmer (58). The nucmer module was used to perform pairwise genome alignment, and single nucleotide polymorphisms (SNPs) and indels were identified from alignment outputs using show-snps.

### 2.11. Statistical analysis

The conjugation frequency among experimental groups were compared using Kruskal-Wallis test with a significant value set at *p* <0.05. Additionally, Dunn’s post hoc test with Holm-adjusted *p*-values was performed for pairwise comparisons. Kaplan–Meier survival analysis was used to evaluate cell persistence over time. Log-rank test was used to assess variations in survival distributions among treatment groups. Kruskal-Wallis and Dunn’s post hoc analyses were performed using *FSA* v0.10.1 package while Kaplan-Meier survival curves were generated using the *survival* v3.8-6 and *survminer* v0.5.2 packages in R v4.2.0.

## 3. RESULTS

### 3.1. Comparative analysis of *bla*_OXA-72_ – harboring plasmid among *Acinetobacter* species

The complete genome of the donor *A. pittii* (EA01) is comprised of 2 plasmid replicons, an 8,502bp mobilizable plasmid (pApiEA01a) which harbors *bla*_OXA-72_ and a 69,083bp conjugative plasmid (pApiEA01b) **(Fig. 1.A.)**. The plasmid mob typing revealed that plasmid pApiEA01a possessed a relaxase and belongs to MOBQ family. A total of 9 coding sequences (CDS) were predicted in pApiEA01a. Notably, toxin-antitoxin (TA) systems and insertions sequence (IS) elements were absent in this plasmid. The *bla*_OXA-72_ gene is flanked between XerC/D regions **(Fig. 1.B.)**. Plasmid rep typing revealed that pApiEA01a harboring *bla*_OXA-72_ belongs to R3-T8. Additionally, the backbone of pApiEA01a shared homology with several plasmids obtained from *Acinetobacter* species, such as pAR3651_1 (CP135161, *A. pittii*, China; identity 99.93%, coverage 67%), pIEC338SCOX (CP015146, *A. pittii*, Brazil; identity 99.96%, coverage 67%), pAba10042a (CP023027, *A. baumannii*, Mexico; identity 99.75%, coverage 66%), pA52-OXA-72 (CP034097, *A. baumannii*, China; identity 94.83%, coverage 26%), and pAbIHIT32296 (KY704308, *A. baumannii*, Luxembourg; identity 95.47%, coverage 13%) **(Fig. 1.C.)**.

**Fig. 1.**
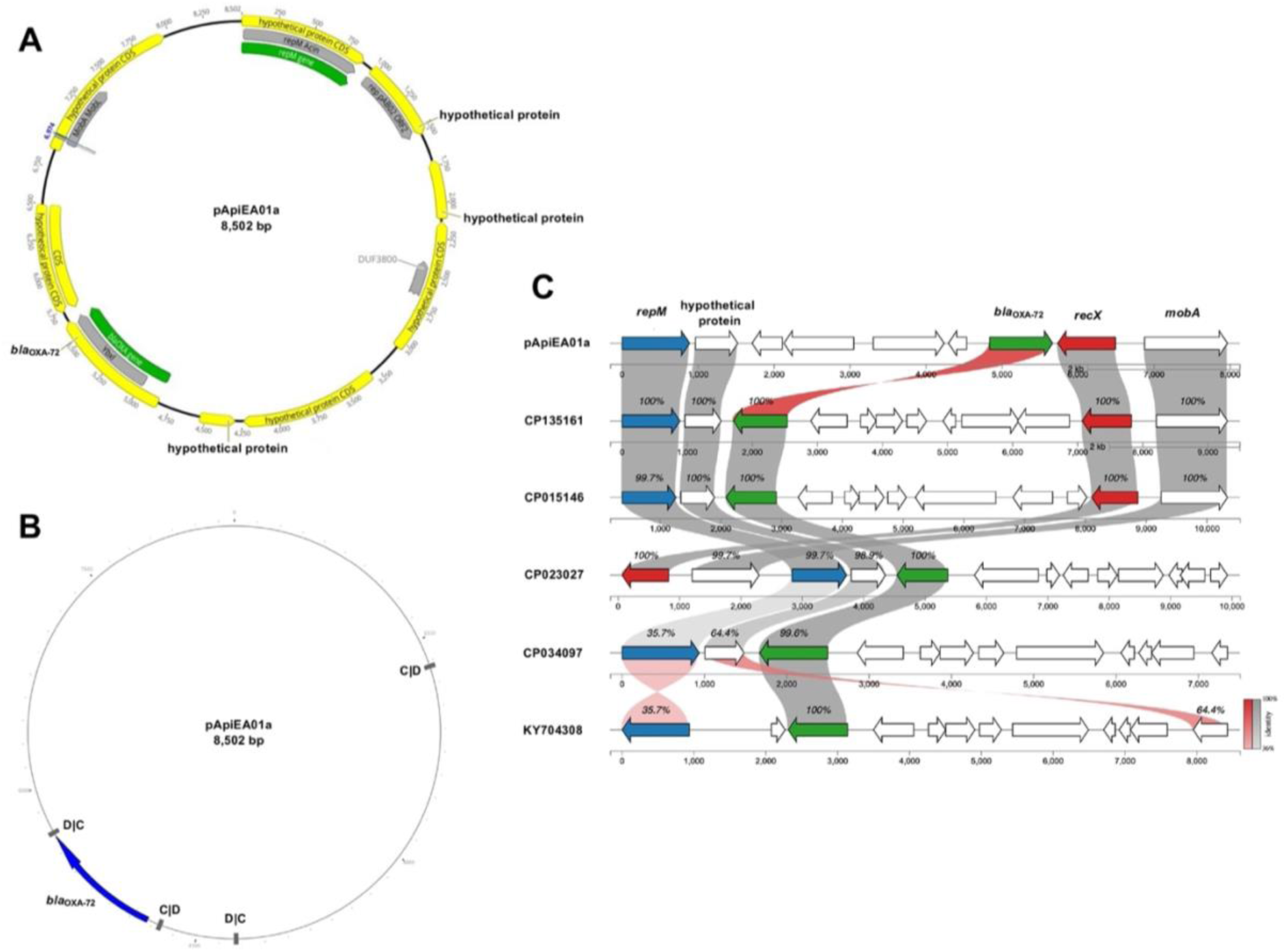
*b/a*_OXA-72_ synteny analysis. **A.** Circular visualization of the mobilizable plasmid pApiEA01a harboring the *bla*_OXA-72_ gene. **B.** Genomic context of *bla*_OXA-72_ gene flanked between XerC/D regions. **C.** Synteny analysis of the *bla*_OXA-72_ – harboring plasmid (pApiEA01a) with other *Acinetobacter* plasmids harboring the *bla*_OXA-72_ gene.

### 3.2. Effect of antibiotic pressure and temperature in conjugation dynamics

We obtained putative transconjugants in one donor-recipient pair, between *A. pittii* and *A. baumannii* **(Fig. 2.A.)**. We showed in this donor-recipient pair that the apparent frequency of putative transconjugants consistent with PRC-detectable marker genes differed according to incubation temperature and presence ot absence of meropenem. **(Fig. 2.B.)**. In setup with and without antibiotic pressure, conjugation was more efficient when donor and recipient were incubated at 27°C compared to 37°C. Additionally, setup with antibiotic showed higher conjugation frequency than setup without antibiotic when incubated at 27°C. Further, conjugation was not detected under 17°C with or without antibiotic pressure. PCR-detectable marker genes such as the plasmid-borne ARG *bla*_OXA-72_ **(Fig. 2.C.)**, plasmid-specific gene *traC* **(Fig. 2.D.),** and chromosomal aminoglycoside resistance gene *strA* **(Fig. 2.E.)** were identified in the putative transconjugants. The 16S rRNA gene sequence confirmed that the putative transconjugants were *A. baumannii* **(Supplementary Table S3)**. The conjugation frequency of the strains across different experimental conditions is listed in **Supplementary Table S4**. No colonies grew on the donor-only or recipient-only control plates after incubation. This confirms that the growth on the dual-selection plates originated from putative transconjugants rather than donor carryover or spontaneous resistant mutants in the recipient.

**Fig. 2.**
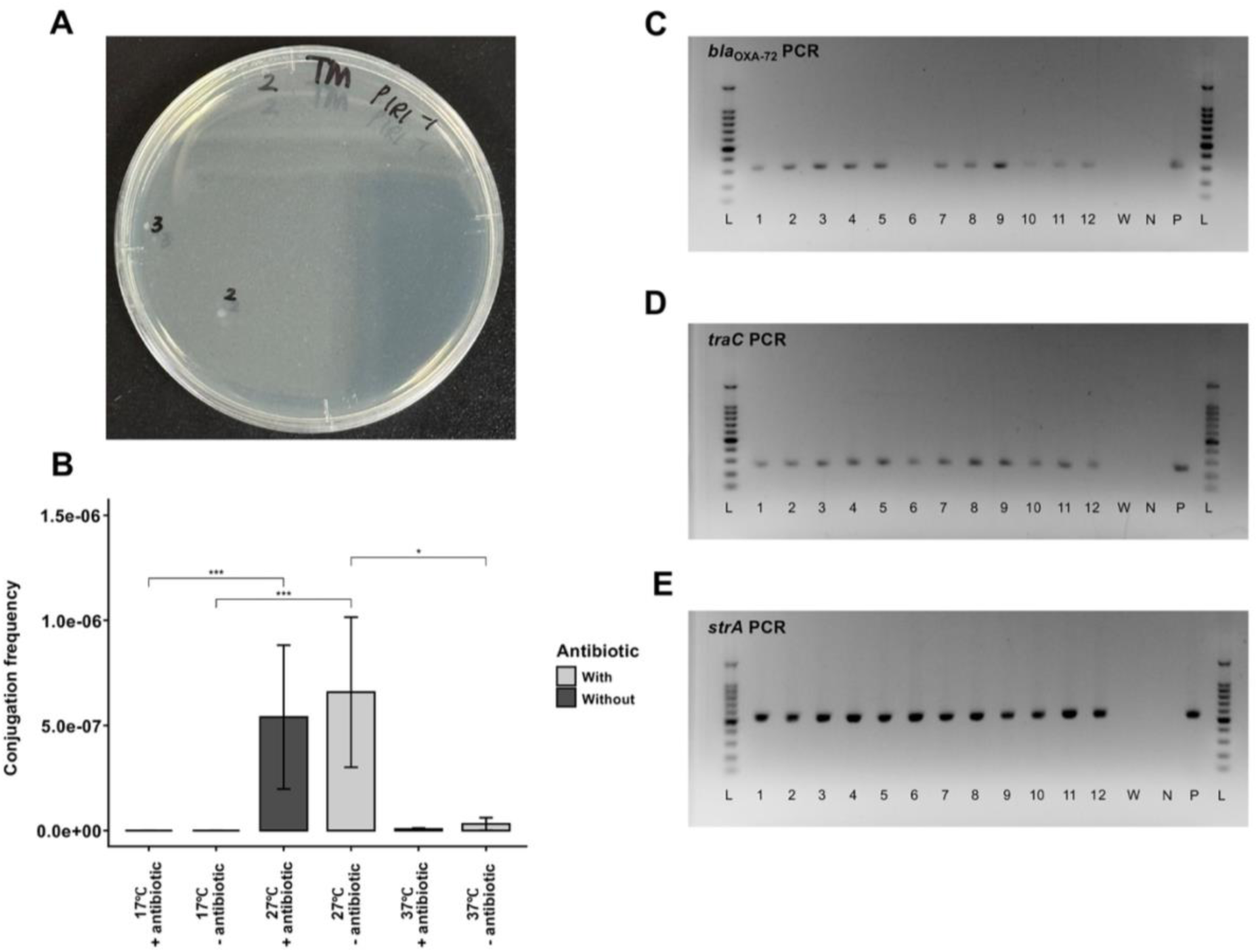
Conjugation assay and PCR-detectable marker gene detection in putative transconjugants. **A.** Putative transconjugants in selective media (tetracycline + meropenem). **B.** Conjugation frequency under varying temperature conditions and antibiotic pressure (\**p* <0.05, \*\*\**p* <0.001). Kruskal-Wallis test was performed to compare conjugation frequency between experimental conditions. **C.** Plasmid ARG *bla*_OXA-72_ in putative transconjugants. **D.** Plasmid specific gene *traC* for conjugative plasmids in putative transconjugants. **E**. Chromosomal ARG *strA* in recipient bacteria (EA06) and putative transconjugant. Positive controls were denoted as “P”: EA01 was used for *bla*_OXA- 72_ and *traC* detection, while EA06 was used for *strA* detection. “N” stands for negative control while “W” means water.

### 3.3. Capsule and outer core oligosaccharide locus typing of donor and recipient *Acinetobacter* strains

We obtained putative transconjugants from only one pair out of the 4 *Acinetobacter* pairs after three repeats. *In silico* typing of capsule (K locus) and outer core oligosaccharide (OC locus) revealed that the donor *A. pittii* EA01 strain was assigned to an unknown KL188 and OC2 locus type, indicating divergence from currently curated reference loci. Among the recipient strains tested, *A. baumannii* EA06, the only strain from which putative transconjugants were obtained, was typed as KL2 and OC1. In contrast, unsuccessful recipients exhibited greater diversity. The clinical *A. baumannii* EC04 strain has a KL9 capsule loci and OC2 outer core loci. While *A. towneri* EA13 strain has a KL179-like capsule loci and OCL6-like outer core loci. Lastly, *A. johnsonii* EA18 strain has a KL237-like capsule loci and OC1-like outer core loci.

Further, investigation of the capsule biosynthesis genes revealed notable differences in the integrity of the *wza–wzb–wzc* operon. The recipient *A. baumannii* strain EA06 which we obtained putative transconjugants harbored *wza*, *wzb*, and *wzc* genes with 100% identity and full-length coverage relative to reference sequences **(Fig. 3.A.)**. In comparison, unsuccessful recipients showed varying degrees of sequence divergence, with *wzb* and *wzc* identities ranging from 99.3% to 99.7% in clinical *A. baumannii* strain EC04 **(Fig. 3.B.)**, while Kaptive failed to assign identity and coverage values for *wza*, *wzb*, and *wzc* genes in *A. towneri* strain EA13 **(Fig. 3.C.)**. Notably, the *wzc* in *A. johnsonii* strain EA18 appeared truncated (77.8% sequence identity and 65.3% coverage) while sequence identity and coverage values were not assigned for *wza* and *wzb* genes **(Fig. 3.D.)**.

**Fig. 3.**
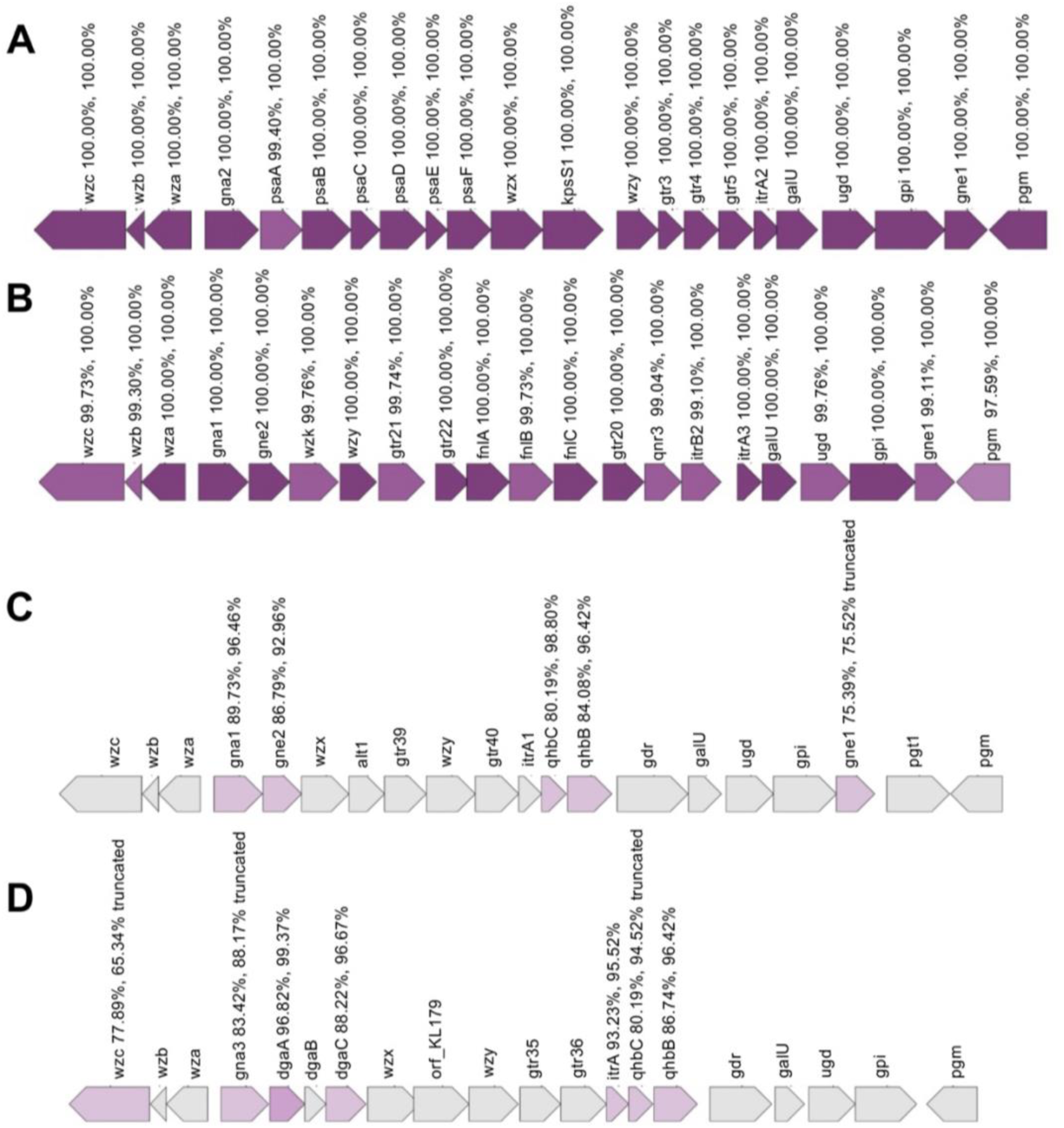
K-locus of the recipient *Acinetobacter* strains. **A.** The strain *A. baumannii* (EA06) from which putative transconjugants were recovered possessed a KL2 K-locus. **B.** The clinical *A. baumannii* (EC04) possessed a KL9 K-locus. **C.** The pig wastewater *A. towneri* has an unknown KL179 K-locus. **D.** The *A. johnsonii* from the pig wastewater has an unknown KL237 K-locus. Gene color intensity indicates the quality of alignment to the reference sequence. Dark shading represents high-confidence alignments with near-complete coverage and high sequence identity, while light shading signifies lower alignment quality caused by reduced coverage, lower identity, or sequence variation. The values shown above each gene (e.g 100% 100%) represent the percent coverage and percent identity of the query gene in relation to the reference.

### 3.4. Defense system frequency and composition in *Acinetobacter* recipient strains

Subsequently, defense systems among the *Acinetobacter* strains were also determined. Putative transfer was observed between two environmental *Acinetobacter* strains (EA01 – *A. pittii* and EA06 – *A. baumannii*) from municipal wastewater. There were no putative transconjugants detected after three repeats from pairs using the recipients *A. baumannii* (EC04) from the hospital, and *A. towneri* and *A. johnsonii* (EA13 and EA18, respectively) from pig farm wastewater. The three recipient *Acinetobacter* strains from which no putative transconjugants were recovered possessed various bacterial defense systems. The clinical *A. baumannii* EC04 strain possessed 6 defense systems, while *A. towneri* EA13 strain and *A. johnsonii* EA18 strain possessed 17 and 12 defense systems, respectively (**Fig. 4.A.)**. Likewise, these recipient strains possessed high frequency of defense genes **(Fig. 4.B.)**. Interestingly, among the defense systems detected in these recipient strains, CRISPR-Cas and anti-plasmid system Wadjet-I were present. EC04 possessed CRISPR-Cas in the chromosome, while EA18 harbored a CRISPR-Cas-bearing plasmid. Additionally, EA13 possessed chromosomal Wadjet-I. Conversely, the strain *A. baumannii* EA06, from which putative transconjugants were obtained, only possessed SspBCDE, PD-T4-5, and PD-T7-5 **(Fig. 4.C.)**. Further analysis showed that the CRISPR-Cas systems are Type I-F system. We found that the CRISPR-Cas spacers showed perfect (100% sequence identity and 0 mismatches) or near-perfect matches (e.g. 94% sequence identity and 1 mismatch) with the conjugative plasmid pApiEA01b **(Supplementary Fig. S2 & S3)**. However, we failed to detect the protospacer adjacent motifs (PAM) for Type I-F in the plasmid sequence.

**Fig. 4.**
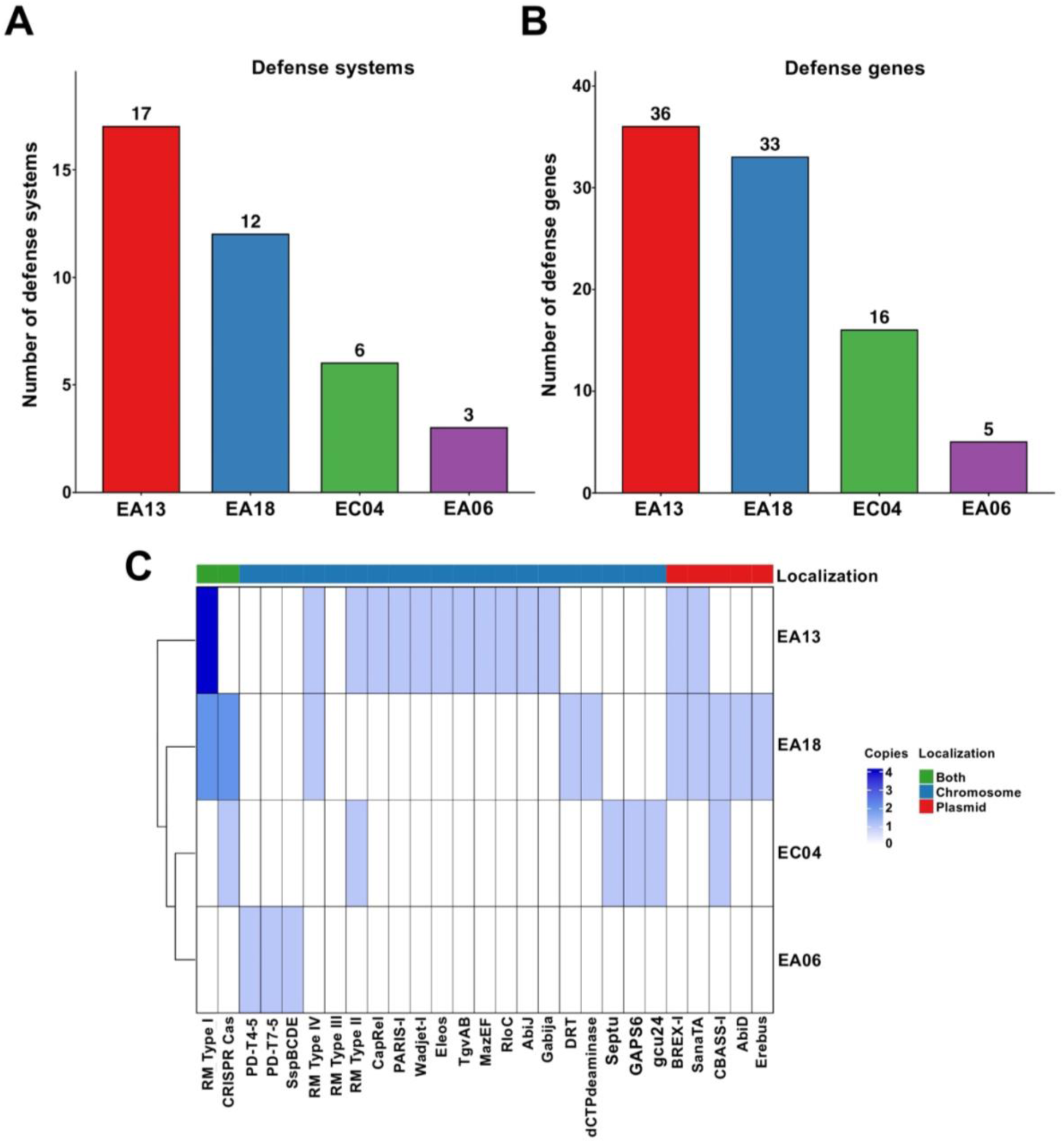
Bacterial defense systems in the recipient *Acinetobacter* strains. **A.** Number of copies of bacterial defense systems among *Acinetobacter* recipient strains. **B.** Number of defense genes among *Acinetobacter* recipient strains **C.** Types of bacterial defense systems among *Acinetobacter* recipient strains.

### 3.5. Phenotypic conversion and fitness cost in putative transconjugant

The putative transconjugant obtained between *A. pittii* and *A. baumannii* exhibited increased meropenem minimum inhibitory concentration and showed fitness cost. Additionally, putative transconjugants exhibited slower growth rate compared to the original recipient bacteria EA06. A prolonged lag phase was observed at the first 4 hours of incubation (2H to 8H) **(Fig. 5.A.)**. We observed a significant difference in terms of growth rate between recipient and putative transconjugants at 8-hour period. Further, the putative transconjugants exhibited an increase in meropenem MIC ranging from 4ug/ml to 8ug/ml which is consistent with *bla*_OXA-72_-associated resistance and supports the PCR-based detection of the gene in these isolates. The MIC of the putative transconjugants was compared to the MIC (2ug/ml) of the original recipient bacteria EA06 **(Fig. 5.B.)**.

**Fig. 5.**
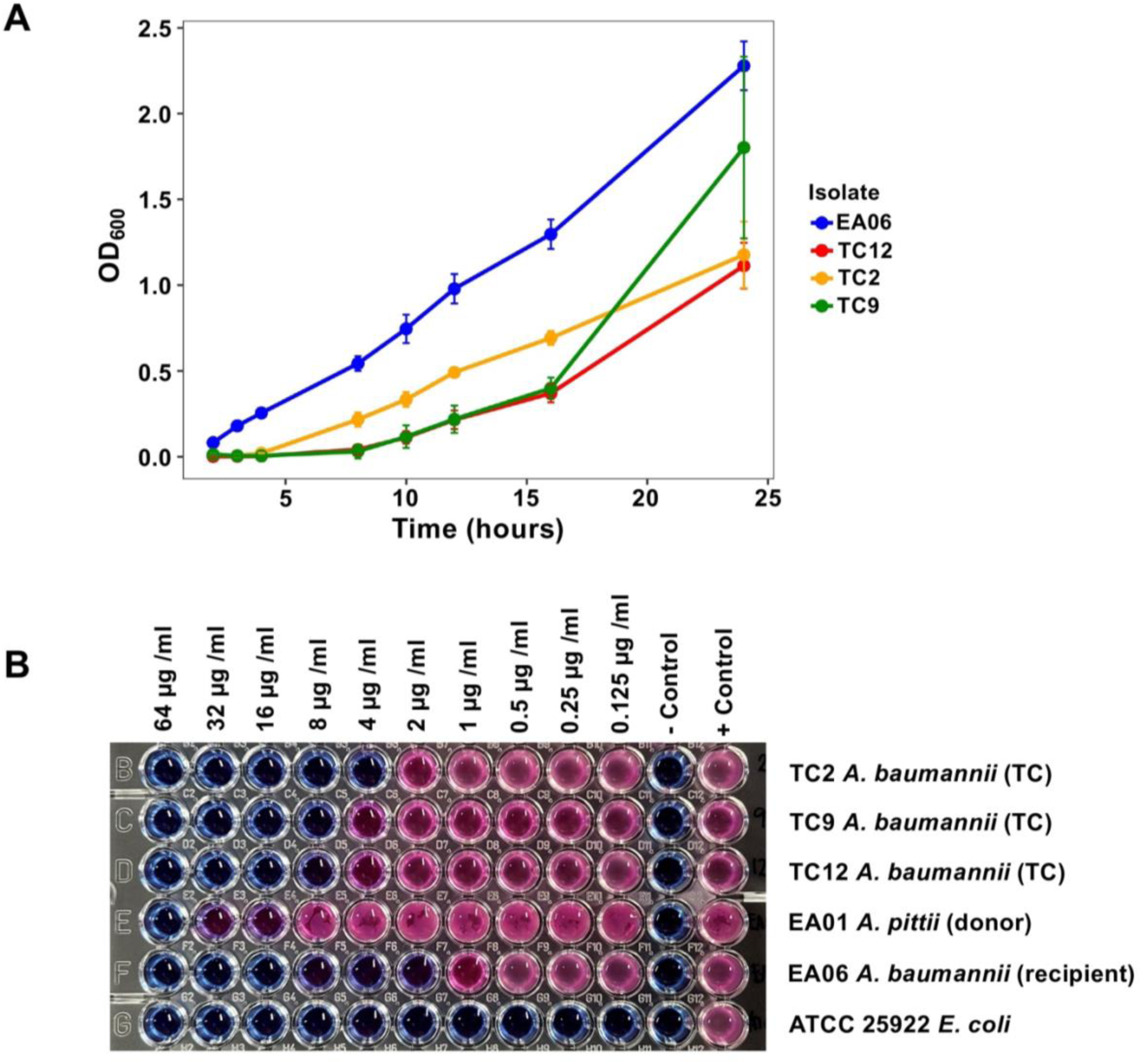
Phenotypic conversion and fitness cost in putative transconjugant. **A.** Growth curve of original recipient (EA06) depicted in blue circles. Slow growth rate of putative transconjugant (TC2, TC9, TC12) is shown in green, red, and purple. Three replicates were used in measuring the OD_600_. Data are means ± SD (*n*=3). **B.** Broth microdilution method showing an increase in meropenem MIC in putative transconjugants (TC). Donor bacteria EA01 and recipient bacteria EA06 were used as the basis for MIC comparison, while *E. coli* ATCC 25922 was used for quality control.

### 3.6. Persistence of putative transconjugants

Kaplan–Meier survival curve of cell persistence at 17°C, 27°C, and 37°C revealed no statistically significant difference among the three different setups. In setup A, cells remained detectable up to day 3 at 17°C; however the difference was not statistically significant (log-rank test, *p* = 0.1) **(Fig. 6.A.)**. Similarly, in setup C, one replicate exhibited slightly prolonged detection of cells at 27°C, but no significant difference was observed (log-rank test, *p* = 0.98) **(Fig. 6.B.)**. PCR screening detected all marker genes (chromosomal ARG *strA*, conjugative gene *traC*, and carbapenem resistance gene *bla*_OXA-72_) in putative transconjugants at varying time points depending on temperature and meropenem concentration. **(Supplementary Fig. S4 – S6)**. Specifically, in the absence of meropenem (setup A), PCR-detectable marker genes were observed up to day 3 at 17°C, day 4 at 27°C, and day 7 at 37°C. Under 2ug/ml meropenem supplementation (setup B), detection was limited to day 1 at 17°C, day 4 at 27°C, and day 3 at 37°C. With 4ug/ml meropenem, marker persisted until day 1 at 17°C, and up to day 6 at both 27°C and 37°C. This may imply that the persisting cells are putative transconjugants consistent with the presence of PCR-detectable marker genes at these specific time points. Interestingly, in the absence of antibiotic pressure (setup A), cells were still recovered until day 30 at 37°C. Nevertheless, cell persistence at day 30 in setup A was not significantly different from other conditions (log-rank test, *p* = 0.09) **(Fig. 6.C.)**. Despite continued growth, plasmid-associated marker genes and the ARG (*traC* and *bla*_OXA-72_) were no longer detected by PCR in day 30 colonies. The 16S rRNA gene sequence confirmed that the persisting cells were *A. baumannii* (**Supplementary Table S5).** Mapping of the Oxford Nanopore reads showed reliable detection of the chromosomal *strA* gene across all isolates, but very few reads mapped to the *bla*_OXA-72_ gene, *traC*, or the complete plasmid sequences of pApiEA01a and pApiEA01b, even at relaxed mapping quality threshold **(Supplementary Table S6)**. The chromosomal *strA* gene mapped consistently well across all isolates and time points, with mean coverage ranging from approximately 100× to over 300×. In contrast, very few reads mapped to *bla*_OXA-72_, *traC*, or the two plasmid sequences. Plasmid-related targets usually had 0 to 89 mapped reads, resulting in very low mean coverage (mostly under 10×). While PCR confirmed the presence of the marker genes in the putative transconjugant colonies, these genes could not be reliably detected in the *de novo* assembled genomes of isolates recovered on days 0, 1, 3, 4, 6, and 30. These results indicate that the maintenance of PCR-detectable markers was unstable over time.

**Fig. 6.**
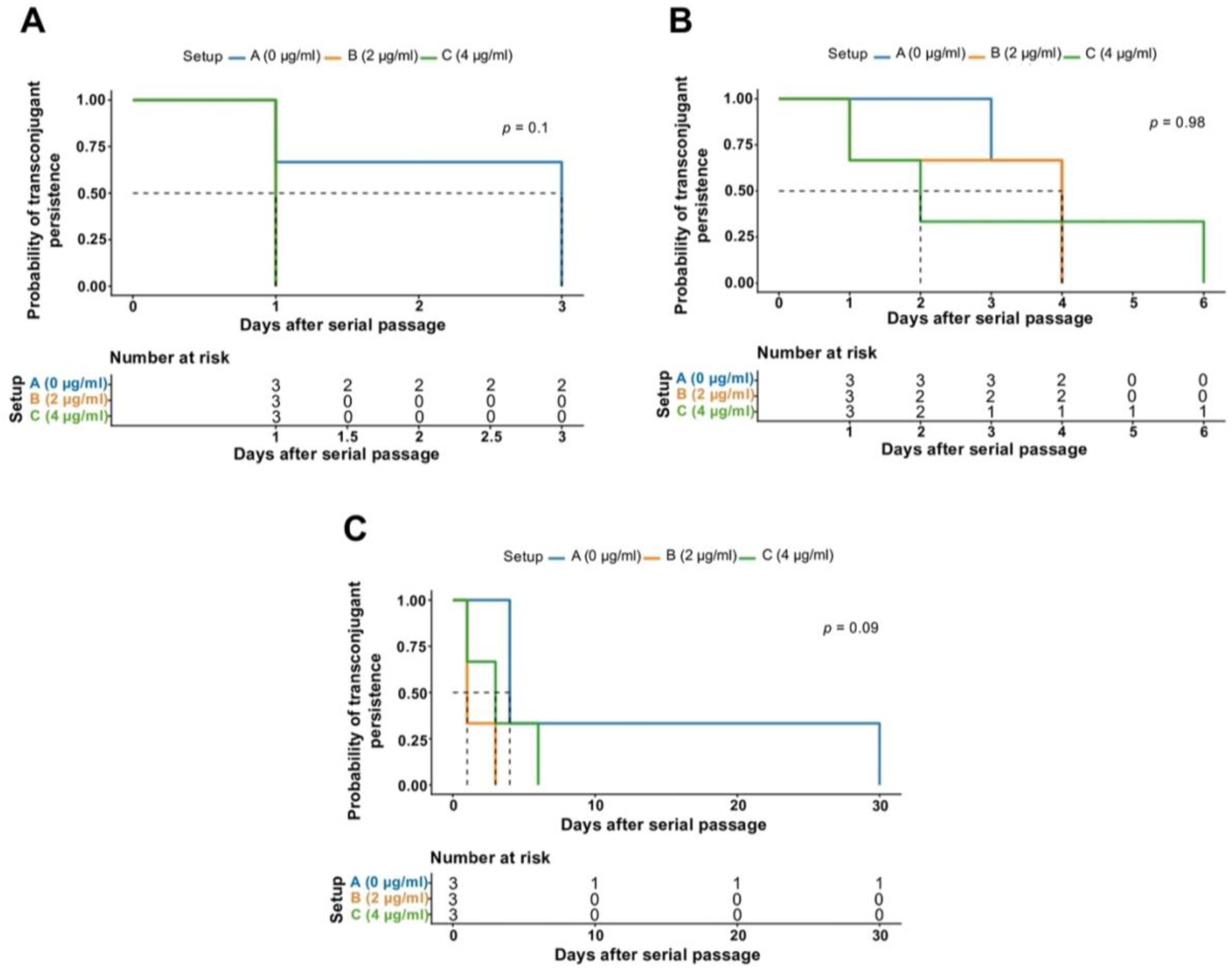
Kaplan–Meier survival curves showing the cell persistence of *Acinetobacter* during 30-day serial passage at different temperatures. **A.** Cell persistence at 17°C under different meropenem concentrations. **B.** Cell persistence at 27°C under different meropenem concentrations. **C.** Cell persistence at 37°C under different meropenem concentrations. Setup A: 0 µg/ml (no meropenem); Setup B: 2 µg/ml meropenem; Setup C: 4 µg/ml meropenem. Curves represent the probability of persisting cells over time. The *p*-value was calculated using the Log-rank test. Risk table indicates the number of replicates with recoverable cells at each time point.

### 3.7. Chromosomal gene mutations in persisting cells at day 30

Comparative genomic analysis of the persisting cells recovered at day 30 identified several genes associated with multidrug efflux, envelope homeostasis, DNA repair, and stress response. We identified an intrinsic carbapenemase gene *bla*_OXA-66_ located at positions 2,287,329–2,288,153 in the chromosome. Additionally, an IS*Aba1* element was detected at positions 2,067,509–2,068,687; however, this element was located approximately 218 kb away from *bla*_OXA-66_ and was therefore not positioned within the immediate upstream regulatory region of the gene.

Interestingly, SNPs and indels were identified in some chromosomal genes **(Table 1)**. The most notable finding was the presence of the carbapenemase gene *ccrA2* (895_896delGA). In addition to carbapenem resistance, multiple multidrug efflux-associated genes were detected, including *abaF, abaQ, amvA, acrA,* and *emrAB*. These genes are associated with active efflux of structurally diverse antimicrobial compounds. Furthermore, we detected the presence of *ampD* gene which is a regulator of β-lactamase expression.

**Table 1.**
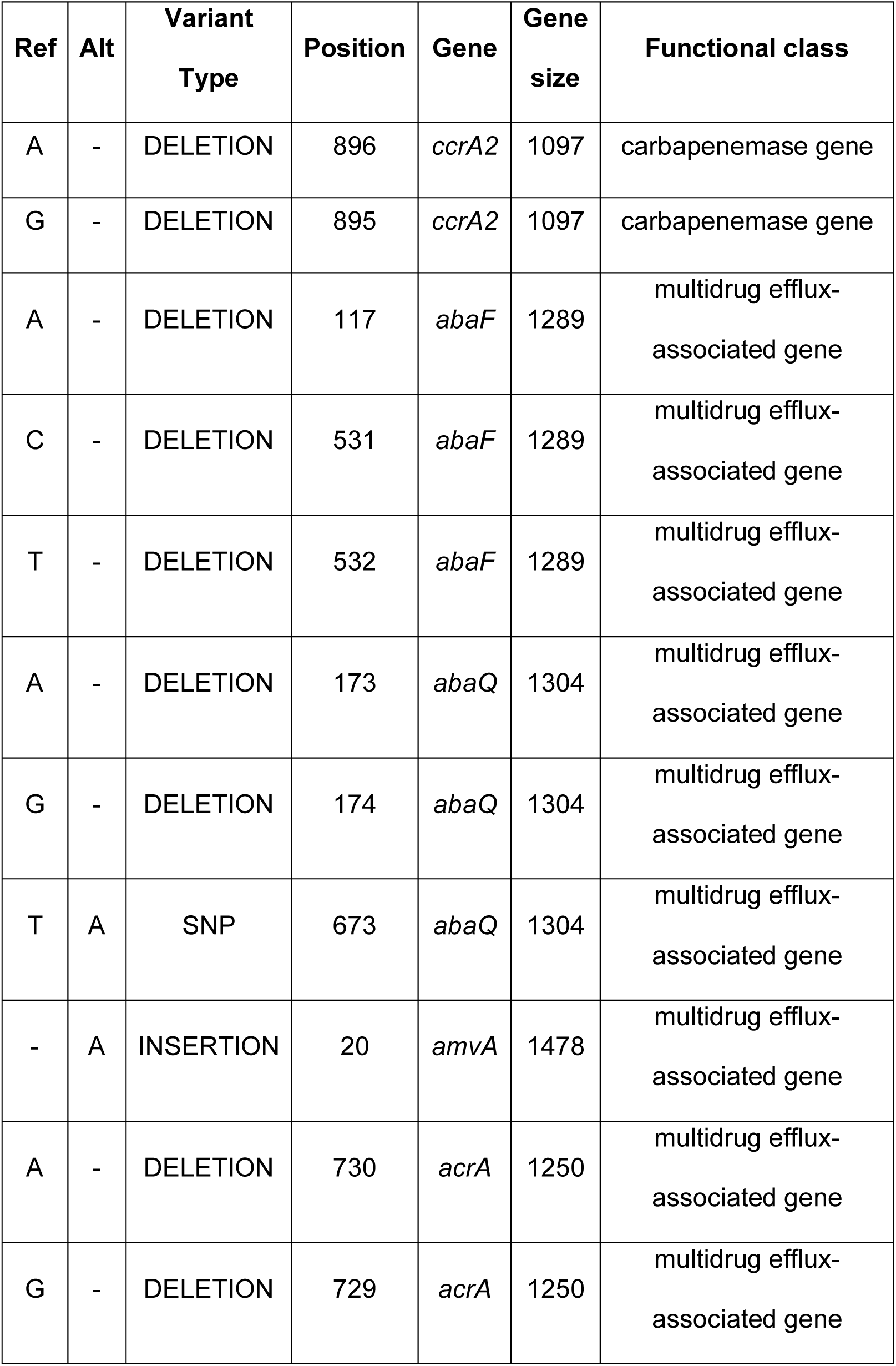
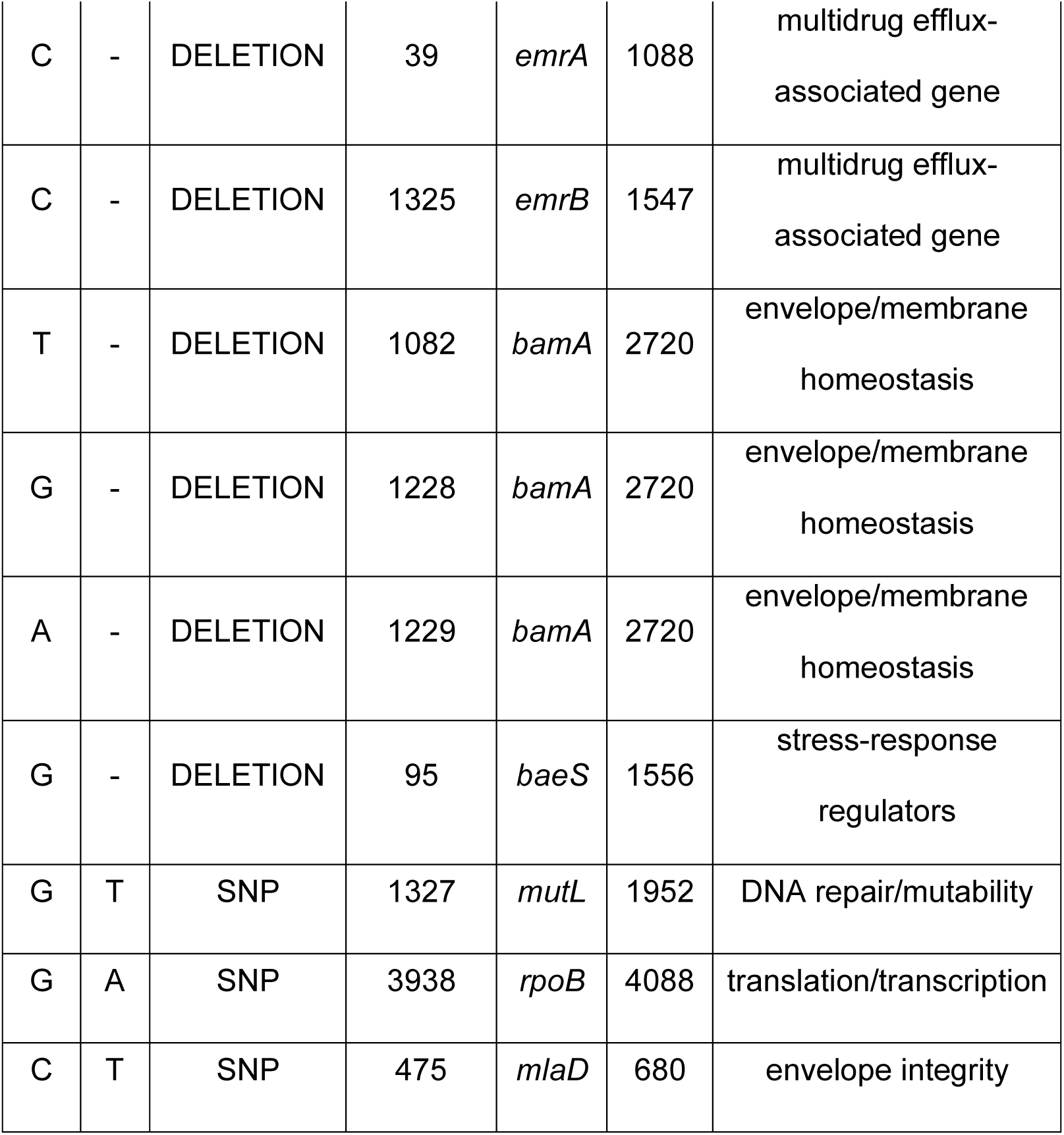
Summary of mutations in chromosomal genes of the persisting cells after 30 days of persistence assay. “Ref” stands for reference allele while “Alt” stands for alternative allele.

## 4. DISCUSSION

### 4.1. Transfer potential of plasmid harboring *bla*_OXA-72_ gene

The emergence of carbapenem resistant *Acinetobacter* species has been reported globally and one of the main mechanisms of resistance is the possession of β-lactamases which are mostly plasmid associated. Thus, it is imperative to probe into the transfer potential and conjugation dynamics of plasmid-borne carbapenemase genes. In this study, we investigated the transfer potential of a plasmid harboring *bla*_OXA-72_ gene from environmental *A. pittii* to other *Acinetobacter* species. Recent studies investigated the transfer potential of plasmid harboring *bla*_OXA-72_ between *Acinetobacter* species (20, 15). He et al. failed to generate transconjugants between clinical *A. pittii* and *A. baumannii* ATCC 17978RIF^R^ due to lack of *oriTs,* T4CPs, and T4SSs; suggesting that the donor *A. pittii* plasmid harboring *bla*_OXA-72_ is deficient to undergo autologous conjugation (20). In contrast, Gheorghe-Barbu et al. successfully demonstrated the conjugation of plasmid harboring *bla*_OXA-72_ from wastewater *A. baumannii* isolates to *A. baylyi* RIF^R^ isolate with a conjugation frequency ranging from 5.4 × 10^-7^ – 3.3 × 10^-13^(15). In this study, PCR-based evidence suggested putative transfer of the *bla*_OXA-72_ gene from *A. pittii* to *A. baumannii*. Detection of *traC* in the putative transconjugants raises the possibility that the conjugative plasmid pApiEA01b may have facilitated the transfer. However, we did not experimentally confirm whether pApiEA01b directly mediated the conjugation. Remarkably, we did not conduct any genetic manipulation or resistance induction in the *Acinetobacter* strains used in the conjugation assay. Consistent with the report of Gheorghe-Barbu et al., we observed low conjugation frequency ranging from 1.5 × 10^-6^ – 9.01 × 10^-9^.

### 4.2. Effects of antibiotic and temperature in conjugation frequency

Subsequently, we observe the effect of varying temperatures and antibiotic pressure in conjugation frequency. One important limitation of this study is that we only recovered putative transconjugants from a single donor-recipient combination. Varying temperatures can impact bacterial growth rates and conjugation efficiency (21, 22, 24–26). Higher temperatures increased cell membrane permeability (23) and transfer frequency by raising the bacterial ATP levels necessary for conjugative transfer (22). Previous study revealed that the optimal temperature for most conjugations to occur is 37°C (25), while conjugation is not observed when cultures are incubated at 4°C (26). Similarly, another study revealed low conjugation frequency of 7.16 × 10^-7^ at 5°C and high at 30°C with a frequency of 2.18 × 10^-5^ (23). While a different study showed that β-lactamase genes in transconjugants were only detected when conjugation assay was carried out at 37°C (24). In contrast with previous studies, we observed higher conjugation frequency under 27°C compared to when incubated under 37°C and 17°C. In this study, both the donor and recipient bacteria were isolated from the influent sample from a wastewater treatment plant with an initial temperature of 26.9°C during sampling. Environmental isolates of *Acinetobacter* typically exhibit optimal growth temperatures around 28-35°C, which is generally lower than the 37°C commonly associated with clinical or host-associated strains (59). This suggests that the temperature conditions used in our study have promoted putative transfer in *A. pittii* and *A. baumannii* pair. However, since putative transconjugants were obtained from only one donor-recipient pair, it remains unclear whether this temperature preference represents a general characteristic of conjugation among *Acinetobacter* isolates.

While it is known that antibiotics serve as a selection pressure in conjugation, the combined effect of antibiotic pressure and varying temperatures in conjugation frequency are still unknown. Previous studies reported that antibiotics enhanced conjugation between bacteria (28–30). In *E. coli*, plasmid transfer increased after exposure of strains in cefotaxime and ampicillin, similarly when also exposed in ciprofloxacin (28). Interestingly, combination of low doses of kanamycin and streptomycin can also promote conjugation in *E. coli* (30). Additionally, gentamicin can also promote conjugation between *E. coli* and *P. aeruginosa* by inhibiting quorum sensing (29). Here, we found that the conjugation frequency varies depending on the presence or absence of low meropenem dosage in relation to varying temperatures. We observed higher conjugation frequency in meropenem-supplemented setup under 27°C compared to setup under 17°C and 37°C. More so, meropenem-supplemented culture shows higher conjugation frequency than meropenem-free culture. While low levels of antibiotic pressure may enhance conjugation frequency, excessive stress particularly under suboptimal temperatures may reduce the donor and recipient cell density, thus limiting opportunities for plasmid transfer. Consequently, the effects of temperature and antibiotic observed should be considered specific to the donor-recipient pair from which putative transconjugants were obtained rather than general characteristics of conjugation in *Acinetobacter*.

### 4.3. Potential barriers of conjugation process

This study also explored the conjugation barriers influencing conjugation outcomes in *Acinetobacter*. Among the four donor–recipient pairs tested, putative transconjugants were not obtained from the three recipient strains derived from clinical and pig wastewater environments, suggesting the presence of intrinsic barriers to plasmid acquisition. In *K. pneumoniae*, presence of capsules can influence the conjugation efficiency by decreasing the acquisition rate of specific conjugative plasmids (60). Additionally, it was also revealed that the donor and recipient serotypes can affect conjugation efficiency (31). To explore possible explanations for the failure of conjugative transfer of pApiEA01a, we examined the envelope composition in *Acinetobacter* recipient strains using *in silico* approaches. The OC1 locus type was present in both successful and unsuccessful recipients, suggesting no clear association observed between OC locus types and conjugation success. Notably, the capsule locus (KL type) and more importantly, the integrity of capsule biosynthesis genes, appeared to show a more meaningful pattern. The successful recipient (*A. baumannii* EA06) had a KL2 capsule type with fully conserved *wza*, *wzb*, and *wzc* genes. These genes encode essential components of the capsule export system, with *wzc* acting as a tyrosine autokinase that controls polysaccharide polymerization and export (61). In contrast, unsuccessful recipients showed sequence divergence or truncation in these genes, especially *wzc* gene. Disruption or alteration of *wzc* has been demonstrated to affect capsule formation and modify surface polysaccharide structure (62). Contrary to the reports of Haudiquet et al. (31), where the absence of capsule is associated with higher conjugation efficiency, no such relationship was evident in the present study. Differences in capsule structure might partly explain the variation we observed in the putative plasmid transfer. However, since putative transfer only occurred in one donor-recipient pair, and many factors such as species-specific traits and genomic variations are likely at play, the link between capsule gene integrity and putative transfer should be viewed as hypothesis-generating rather than conclusive.

Beyond bacteria surface-associated factors, this study highlights the possible role of bacterial defense systems as barriers to horizontal gene transfer via conjugation in *Acinetobacter*. Bacterial defense systems, particularly CRISPR-Cas and restriction–modification (RM) systems, are among the most prevalent defense systems limiting horizontal gene transfer (HGT) (63). These systems recognize and neutralize incoming foreign DNA elements, such as plasmids. Consistent with this, multiple defense systems were identified across the recipient *Acinetobacter* strains. Notably, isolates that failed to undergo conjugation harbored a greater number and diversity of defense systems. The clinical strain EC04 contained 6 defense systems, while pig wastewater strains EA13 and EA18 possessed 17 and 12 systems, respectively. In comparison, the successful recipient EA06 displayed a comparatively limited number of defense systems. This pattern is consistent with the possibility that greater number of defense systems may reduce the likelihood of plasmid acquisition, as suggested by previous genomic studies (64). Among the detected defense systems, the presence of CRISPR-Cas systems in strains EC04 and EA18 is particularly noteworthy. These systems provide adaptive immunity by recognizing and degrading foreign nucleic acids in a sequence-specific manner (54, 65). CRISPR-Cas systems are predominantly carried by *Acinetobacter*, some with exceptionally large CRISPR array (66). Additionally, strain EA13 possessed the Wadjet-I anti-plasmid defense system, a recently described mechanism known to restrict plasmid maintenance (64). The presence of Wadjet-I in this isolate raises the possibility that it could also be one of the several factors potentially influencing the absence of putative transconjugants. However, this association remains correlative rather than causal. Additionally, our findings rely on *in silico* predictions of defense systems, so functional assays and larger datasets will be necessary to clarify the exact mechanisms underlying these barriers to HGT.

### 4.4. Phenotypic effects in putative transconjugants

In addition to conjugation frequency, we investigated growth dynamics and fitness cost in the putative transconjugant. The acquisition of new plasmids entails fitness cost, which can result in the disruption of cellular pathways and regulations (67) and alteration of bacterial phenotypes (68). Transconjugants grew slower compared to plasmid-free lineages, showing prolonged lag time (69). Consistent with these findings, the putative transconjugants in this study exhibited reduced growth rates compared to the original recipient strain. The prolonged lag phase observed suggest an adaptive response where the recipient bacteria make metabolic and physiological adjustments associated with the presence of PCR-detectable bla_OXA-72_ marker gene. Despite the apparent fitness cost, the putative transfer of *bla*_OXA-72_ gene increased the meropenem MIC of putative transconjugants by 2-fold concentration compared to the original recipient bacteria. Even slight increases in MIC values can lead to decreased treatment effectiveness and promote the emergence of resistant strains in clinical settings. The increased meropenem MIC is consistent with the putative transfer of carbapenem resistance, as indicated by the PCR detection of *bla*_OXA-72_ gene. However, since the WGS data failed to robustly confirm the presence of the plasmid, these findings need to be interpreted with caution. Taken together, these results highlight that while the presence of PCR-detectable *bla*_OXA-72_ gene incurs a fitness cost, it concurrently enhances carbapenem resistance in the putative transconjugants.

### 4.5. Effects of temperature and antibiotics on cell and marker gene persistence

Despite studies on the conjugative transfer of carbapenemase genes, the conjugation-mediated persistence of the newly acquired plasmid and ARGs in the transconjugant cells under varying antibiotic and temperature conditions remained unclear. In evolution experiments, the decrease in frequency of plasmid hosts incubated under 20°C compared to 37°C was slower, indicating lower plasmid loss in the populations incubated under 20°C (35). While continuous antibiotic supplementation can significantly increase the persistence of plasmid in carrier bacteria (34). In this study, we observed no significant difference in terms of cell persistence under meropenem supplementation and meropenem-free conditions. However, we observed that cultures at 37°C with 4ug/ml meropenem supplementation persisted until day 6, while meropenem-free cultures persisted until day 30. Despite the continued detection of persistent cells throughout the study, the proportion of PCR-marker-positive colonies and *bla*_OXA-72_ gene showed a gradual decline during the 30-day serial passage as confirmed by the absence or presence of PCR-detectable marker genes. Interestingly, Abe et al. (36) reported chromosomal integration of carbapenemase gene in *E. coli* in the absence of antibiotic pressure. In this study, we failed to observe chromosomal integration of *bla*_OXA-72_, which is consistent with the absence of meropenem selection during culture. This could be attributed to the lack of necessary IS elements required for efficient chromosomal integration of *bla*_OXA-72_. In the absence of antibiotic pressure, persistence of resistance is enabled by several factors, including genetically linked resistance genes and other mutations (27). We identified several mutations in certain genes associated with multidrug efflux-associated genes, envelope and membrane genes, and stress-response regulators. These genomic changes were observed in cells that continued to grow on meropenem-supplemented media despite the absence of *bla*_OXA-72_. However, their potential contribution to meropenem tolerance was not functionally validated in this study. Collectively, these findings show that PCR-detectable marker genes in the putative transconjugants were unstable over time. Although we examined the effects of different temperatures and antibiotic concentrations, Kaplan-Meier analysis did not reveal significant differences across these conditions. The gradual decline in the proportion of marker-positive colonies over time suggests that the persistence of *bla*_OXA-72-_associated markers was quite variable. Additionally, the WGS data did not reliably confirm the presence of *bla*_OXA-72_ or confirm plasmid configuration. These observations indicate that long-term persistence of the acquired resistance depend on the interplay between host fitness, environmental adaptation, and selective pressure. Understanding these factors is essential for predicting the long-term dissemination and maintenance of carbapenem resistance genes in bacterial populations.

Lastly, we observed the presence of PCR-detectable marker genes such as *bla*_OXA-72_, *strA*, and *traC* in the putative transconjugants and persisting cells using the conventional PCR. Whereas the *de novo* assembled genome of representative putative transconjugants and persisting cells failed to detect the marker genes *bla*_OXA-72_ and *traC.* Mapping of the Oxford Nanopore reads showed that chromosomal *strA* gene was consistently detected, but only a few reads were mapped to *bla*_OXA-72_, *traC,* and plasmid sequences. Thus, neither the presence of *bla*_OXA-72_ nor the stable retention of putative plasmids in these isolates were supported by the WGS data. As a result, rather than being confirmed transconjugants, these isolates should be considered putative transconjugants based on the presence of PCR-detectable marker genes. In bacterial genomics, this discrepancy between PCR and WGS data is a recognized challenge, especially for plasmid-borne ARGs. This discrepancy between PCR-positive and WGS-negative was reported in a prior study, which revealed false-negative results attributed to fluctuating plasmid copy number and instability during culture (70). Additionally, due to biases introduced during library preparation, long-read sequencing alone often produced assemblies that are structurally complete but occasionally missed plasmids (71). Similarly, long-read-only assemblers most frequently overlooked small plasmids (<10 kb) (72).

This study identifies several limitations that should be recognized when interpreting the findings. The primary limitation of this study is that only one donor-recipient *Acinetobacter* pair yielded putative transconjugants and that the presence and configuration of the associated plasmids in these isolates could not be robustly resolved by WGS. Although PCR detected the marker genes, the inconsistency between PCR and WGS results weakens our conclusions about successful conjugation and plasmid persistence. Additionally, the observed effects of antibiotic and temperature should be interpreted with caution, as they likely reflect pair-specific responses rather than general features of conjugation across *Acinetobacter* species. Another limitation is the lack of experimental validation of bacterial defense systems. While genomic analysis identified the presence of defense-related genes, their functional activity during conjugation was not assessed. Therefore, the observed lack of conjugation in certain *Acinetobacter* strains could be associated with these systems, but a causal relationship cannot be established. Another limitation of this study is that capsule locus and OC locus characterization was conducted only through *in silico* analysis without experimental validation. While possible associations with conjugation efficiency were observed, the direct role of these loci in plasmid transfer and persistence was not functionally confirmed. Additionally, the conjugation efficiencies reported here were derived under controlled laboratory conditions and may not fully reflect the complexity of microbial communities in clinical or environmental settings. As such, the findings should be interpreted as an indication of the physiological potential of these organisms rather than a direct measure of horizontal gene transfer in natural ecosystems. Lastly, the hybrid assembly strategies would likely resolve such plasmid-related discrepancies more reliably in subsequent studies.

## 5. CONCLUSION

To our knowledge, this study provides PCR-based evidence consistent with putative *bla*_OXA-72_ transfer in a single *A. pittii-A. baumannii* donor-recipient pair, although whole genome sequencing data did not robustly resolve the plasmid presence or configuration in the putative transconjugants. Within this pair, antibiotic pressure and temperature were investigated as factors potentially influencing conjugation frequency, but no significant effects on the persistence of the detected marker genes observed. Overall, our findings should be considered exploratory and provide a basis for future studies aimed at clarifying the mechanisms underlying the transfer and persistence of *bla*_OXA-72_ in *Acinetobacter* species.

## Data availability

The complete genome assembly of the putative transconjugants in this study have been deposited in the GenBank database under BioProject accession number PRJNA1480103.

## ACKNOWLEDGEMENTS

We appreciate the help of Arata Danyoshi and Aya Kadoya for the technical assistance in identifying suitable *Acinetobacter* pairs and carrying out the conjugation assays. We are also grateful to Khristina Judan Cruz, Dewi Gustari, Ngure Kagia, and Rina Sumiokada, for their assistance with the persistence assays. This research was performed by the Environment Research and Technology Development Fund (JPMEERF25S21213) of the Environmental Restoration and Conservation Agency provided by Ministry of the Environment of Japan.

**Kozo Watanabe:** is supported by the Environment Research and Technology Development Fund (JPMEERF25S21213) of the Environmental Restoration and Conservation Agency provided by Ministry of the Environment of Japan.

**Kenneth A. Bongulto:** Conceptualization, Methodology, Investigation, Formal analysis, Writing- Original draft, Project administration, Visualization.

**Hisamichi Tauchi:** Methodology, Writing- Reviewing and Editing.

**Satoru Suzuki:** Conceptualization, Methodology, Writing- Reviewing and Editing, Supervision

**Kozo Watanabe:** Conceptualization, Methodology, Formal analysis, Writing- Reviewing and Editing, Funding Acquisition, Project administration, Supervision.

## Supplemental Material

**Supplementary Figure S1. Schematic diagram of the filter-mating assay.**

**Supplementary Figure S2. Analysis of the CRISPR-Cas spacers in EC-4 strain (*A. baumannii*) and conjugative plasmid (pApiEA01b) sequence.**

**Supplementary Figure S3. Analysis of the CRISPR-Cas spacers in EA-18 strain (*A. johnsonii*) and conjugative plasmid (pApiEA01b) sequence.**

**Supplementary Figure S4. Amplification of the chromosomal *strA* gene in the persisting cells.**

**Supplementary Figure S5. Amplification of the plasmid-borne ARG *bla*_OXA-72_ in the persisting cells.**

**Supplementary Figure S6. Amplification of the plasmid marker gene *traC* in the persisting cells.**

**Supplementary Table S1. Genomic characteristics of donor and recipient *Acinetobacter* strains.**

**Supplementary Table S2. Primers and PCR conditions used for the detection of marker genes.**

**Supplementary Table S3. 16S rRNA gene analysis of putative transconjugants.**

**Supplementary Table S4. Colony forming unit of putative transconjugants after selection in LB with meropenem and tetracycline supplementation.**

**Supplementary Table S5. 16S rRNA gene analysis (reverse only) of persisting bacterial cells.**

**Supplementary Table S6. Summary of mapped reads from persistence assay.**

